# A Cytoplasmic GOLD Protein Controls Cell Polarity

**DOI:** 10.1101/379057

**Authors:** Deike J. Omnus, Angela Cadou, Gary H.C. Chung, Jakob M. Bader, Christopher J. Stefan

## Abstract

Phosphoinositide lipids provide spatial landmarks during polarized secretion. Here, we elucidate a role for phosphatidylinositol 4-phosphate (PI4P) metabolism in the control of cell polarity. In budding yeast, PI4P is enriched at the plasma membrane of growing daughter cells. Upon heat shock however, PI4P rapidly increases at the plasma membrane in mother cells resulting in a more uniform PI4P distribution. Rather than phosphoinositide kinase activation, PI4P hydrolysis is impaired to generate the heat-induced PI4P signal in mother cells. This fine tune control of PI4P metabolism is mediated through attenuation of the Osh3 protein that binds and presents PI4P to a phosphoinositide phosphatase. Importantly, Osh3 undergoes phase transitions upon environmental stress conditions, resulting in intracellular aggregates and reduced cortical localization. The chaperone Hsp104 co-assembles with intracellular Osh3 granules, but is not required for their formation. Interestingly, the Osh3 GOLD domain, also present in the ER-localized p24 cargo adaptor family, is sufficient to form stress granules. Accordingly, GOLD-mediated phase transitions may provide a general mechanism to modulate secretion and growth upon transient changes in physiological and environmental conditions.

## Introduction

Phosphatidylinositol 4-phosphate (PI4P) is emerging as a key determinant of plasma membrane (PM) identity and function. Previous work indicates vital roles for PI4P in the control of PM ion channels and the general recruitment of polybasic proteins to the PM (Hammond *et al*., 2012). PI4P may even serve as a spatial cue or signpost for protein targeting to specialized PM domains. For instance, PI4P is highly enriched in the primary cilium of neural progenitor cells (Chavez *et al*., 2015; Garcia-Gonzalo *et al*., 2015). In budding yeast, PI4P organizes the actin cytoskeleton and targets the p21-activated kinase Cla4 to sites of polarized growth (Audhya *et al*., 2000; Wild *et al*., 2004). Accordingly, defects in PI4P metabolism impair polarized secretion in yeast (Kozminski *et al*., 2006). However, regulatory mechanisms that control PI4P distribution at the PM have not been extensively characterized.

Synthesis of PI4P at the PM is carried out by phosphatidylinositol 4-kinase type IIIα (PI4KIIIα encoded by PI4KA in mammals and the *STT4* gene in *Saccharomyces cerevisiae*) (Audhya *et al*., 2000; Balla *et al*., 2005; Nakatsu *et al*., 2012). The yeast Stt4 PI4K localizes to cortical assemblies termed PIK patches, consistent with its essential role in generation of PI4P at the PM (Audhya *et al*., 2000; Baird *et al*., 2008). Curiously however, while PI4P is enriched in the PM of growing daughter cells (buds), cortical assemblies of the Stt4 PI4K are found extensively in mother cells (Audhya and Emr, 2002; Baird *et al*., 2008). It is unknown how PI4P accumulates in the growing bud and how PI4P levels are kept relatively low in mother cells where Stt4 PIK patches reside. Here, we address this paradoxical distribution between the Stt4 PI4K and its product PI4P. We find that the Stt4 PI4K is associated with the cortical endoplasmic reticulum (ER) in mother cells. Junctions between the ER and PM, termed ER-PM contacts, are sites for PI4P-mediated lipid exchange reactions carried out by the conserved ORP/Osh lipid transfer proteins (Chung *et al*., 2015b; Moser von Filseck *et al*., 2015; Sohn *et al*., 2016). Accordingly, we find that non-vesicular lipid exchange taking place at ER-PM junctions controls the level and distribution of PI4P at the PM.

Importantly, we show that external stimuli such as changes in environmental conditions influence PI4P utilization. Upon heat shock conditions known to disrupt polarized secretion (Delley and Hall, 1999), non-vesicular PI4P consumption is attenuated, resulting in the generation a PI4P signal in mother cells and loss of PI4P polarity. This control is achieved through inactivation of the Osh3 lipid transfer protein that normally binds and delivers PI4P to an ER-localized PI4P phosphatase. Interestingly, Osh3 is a stress-sensitive protein and rapidly forms intracellular aggregates upon heat shock. Notably, localization of the human ORP5 protein is dynamically regulated in response to calcium signals. We propose that the control of PI4P catabolism provides a conserved mechanism to direct polarized growth and cellular responses to extracellular signals.

## Results

### PI4P Metabolism Controls Its Distribution

Phosphoinositide lipid metabolism at the plasma membrane (PM) is modulated by specific lipid kinases and phosphatases (Fig 1A). Yet how cells generate discrete phosphoinositide signals at the PM in response to physiological stimuli is still poorly understood. The phosphoinositide isoform phosphatidylinositol 4-phosphate (PI4P) may play an especially important, but underappreciated, role in polarized cell growth and cell signaling. Previous studies have reported that PI4P is enriched in small (growing) daughter cells in the budding yeast (Stefan *et al*., 2011; Manford *et al*., 2012; Omnus *et al*., 2016). However, these studies described qualitative observations rather than quantitative measurements. We monitored PI4P levels and localization at the PM by quantitative microscopy (Figs 1 and EV1), using a validated PI4P FLARE (fluorescent lipid-associated reporter), the P4C domain of SidC that specifically binds PI4P (Luo *et al*., 2015). Because PI4P localizes to intracellular compartments as well as the PM, GFP-P4C intensities co-localized with the PM marker (mCherry-2xPH^PLCδ^) were specifically measured (Fig EV1A). Under nonstress (26°C) growth conditions, the PI4P FLARE was highly enriched at the PM of daughter cells compared to mother cells (> 7-fold, Figs 1B-C; see examples in EV1A-B). This polarized distribution suggests that PI4P may serve as a landmark at the PM, as has been proposed for other negatively charged lipids including PI(4,5)P_2_ and phosphatidylserine (He *et al*., 2007; Garrenton *et al*., 2010; Fairn *et al*., 2011; Hammond *et al*., 2012).

**Figure 1.**
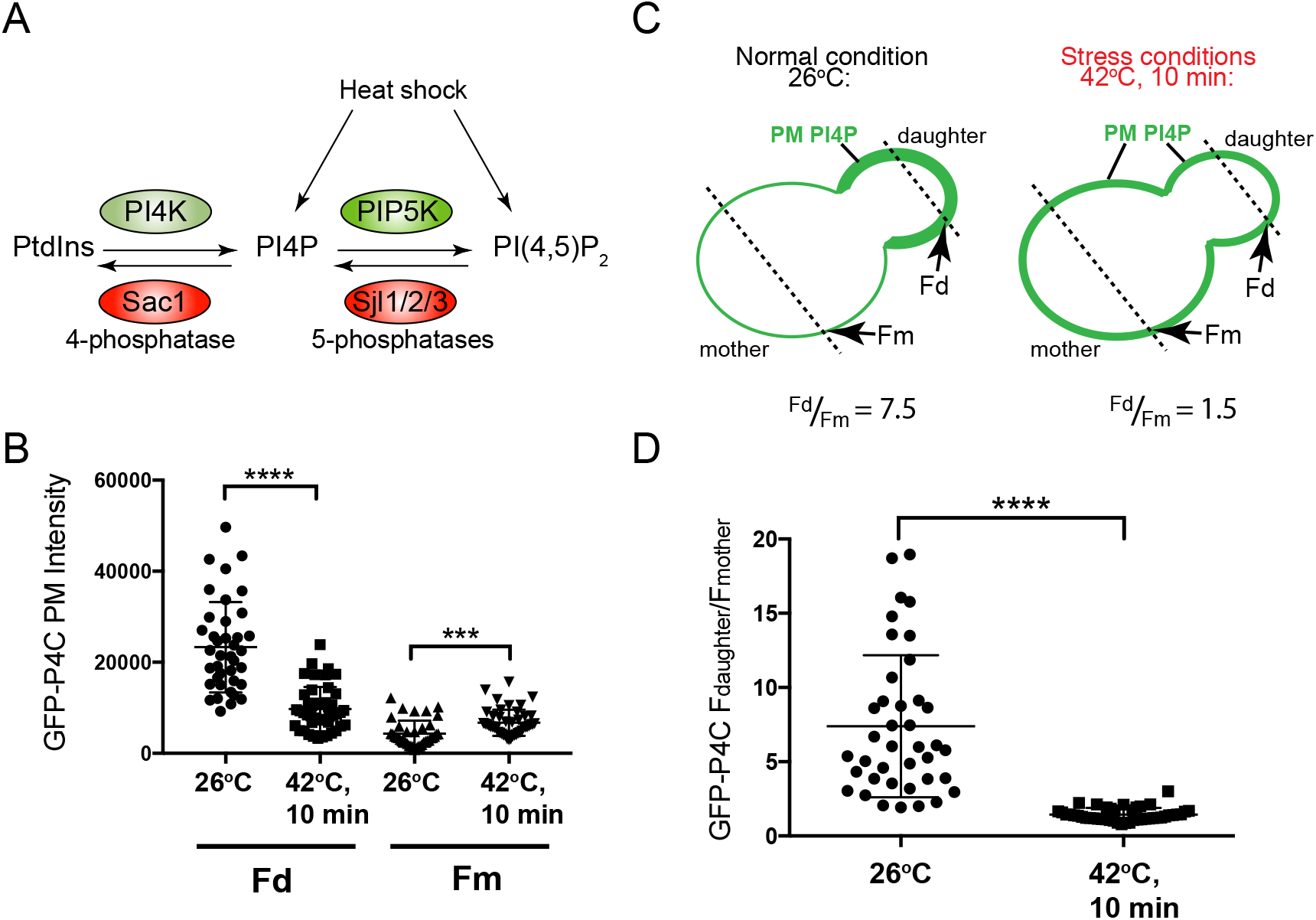
PI4P distribution is regulated by growth conditions. **(A)** Schematic representation of kinases and phosphatases involved in PI4P and PI(4,5)P_2_ metabolism. Heat shock leads to elevated levels of both PI4P and PI(4,5)P_2_. **(B)** Quantitation of GFP-P4C fluorescence at the plasma membrane (PM) of daughter (Fd) and mother (Fm) cells at 26°C and after a 10 min 42°C heat shock. Graph shows the Fd and Fm mean values of individual cells. Total number of cells analyzed in three independent experiments: 26°C, n=40, 10min 42°C, n=45. Error bars show standard deviations. The changes in Fd and Fm after heat shock are statistically significant (t test, ****p< 0.0001, ***p=0.0002). **(C)** Schematic representation of the method used to measure PM GFP-P4C fluorescence intensities at 26°C and after 42°C heat shock. Briefly, line scans were applied through both daughter and mother cells using Fiji and the peak values corresponding to the GFP-P4C fluorescence intensity at the PM in the daughter (Fd) and mother cell (Fm) were recorded to calculate Fd/Fm ratios. **(D)** Quantitation of GFP-P4C fluorescence at the plasma membrane (PM) of daughter (Fd) and mother (Fm) cells at 26°C and after a 10 min heat shock as described in C. Graph shows the Fd/Fm ratio of individual cells. Total number of cells analyzed in three independent experiments: 26°C, n=40, 10min 42°C, n= 45. Error bars show standard deviations. The decrease in the Fd/Fm ratio after heat shock is statistically significant (t test, ****p< 0.0001). Scale bars, 5 μm.

If PI4P serves as a spatial landmark at the PM, then its distribution should change in response to physiological stimuli that regulate cell polarity. Heat elicits several responses in yeast cells including reorganization of the actin cytoskeleton, Rho GTPase signaling, and increased vesicle trafficking events in mother cells (Delley and Hall, 1999; Audhya and Emr, 2002; Zhao *et al*., 2013). Upon heat shock (10 minutes at 42°C), the polarized enrichment of PI4P in daughter cells was significantly reduced. There was a two-fold decrease in GFP-P4C intensity at the PM in daughter cells (Figs 1B, EV1C) and a two-fold increase at the PM in mother cells (Figs 1B, EV1D). These responses occurred rapidly within 2-4 minutes (Figs EV1B-D). Consequently, the ratio of PI4P distribution between daughter and mother cells decreased significantly (5-fold) following heat shock (F_d_/F_m_=7.5 at 26°C versus F_d_/F_m_ =1.5 at 42°C; Fig 1D).

Next we investigated potential mechanisms for the increase in PI4P at the PM in mother cells, which could result from either increased synthesis or decreased hydrolysis. PI4P is generated at the PM in yeast by the Stt4 protein, an ortholog of phosphatidylinositol 4-kinase type IĲα (PI4KIIIα) (Audhya *et al*., 2000). Accordingly, GFP-P4C is lost from the PM in temperature conditional *stt4* mutant cells following heat shock (Fig EV2A). PI4KIIIα is present in two distinct protein complexes. PI4KIIIα complex I is comprised of Stt4/Efr3/Ypp1 in yeast and PI4KIIIα/EFR3/TTC7/FAM126A in mammalian cells (Fig 2A) (Baird *et al*., 2008; Nakatsu *et al*., 2012; Baskin *et al*., 2016). PI4KIIIα complex II consists of Stt4/Efr3/Sfk1 in yeast and PI4KIIIα/EFR3/TMEM150 in mammalian cells (Fig 2A) (Audhya and Emr, 2002; Chung *et al*., 2015a). Sfk1 is necessary for heat-induced PI(4,5)P_2_ synthesis in yeast and the TMEM150 proteins are involved in PI(4,5)P_2_ re-synthesis following phospholipase C activation (Audhya and Emr, 2002; Chung *et al*., 2015a). We tested if the heat-induced increase in PI4P at the PM in mother cells required the ‘signaling’ Stt4 PI4K complex II implicated in heat-induced PI(4,5)P_2_ synthesis. Surprisingly, heat shock induced a PI4P signal at the PM in mother cells lacking Sfk1 (Fig 2B). Thus, the Sfk1-containing Stt4 PI4K complex II is not required for heat-induced PI4P redistribution from daughter cells to mother cells. Instead, the Stt4 PI4K complex I may generate the inducible PI4P signal in mother cells. In support of our results, a recent study has implicated PI4K complex I in stimulus-induced PI(4,5)P_2_ synthesis in metazoan cells (Balakrishnan *et al*., 2018).

**Figure 2.**
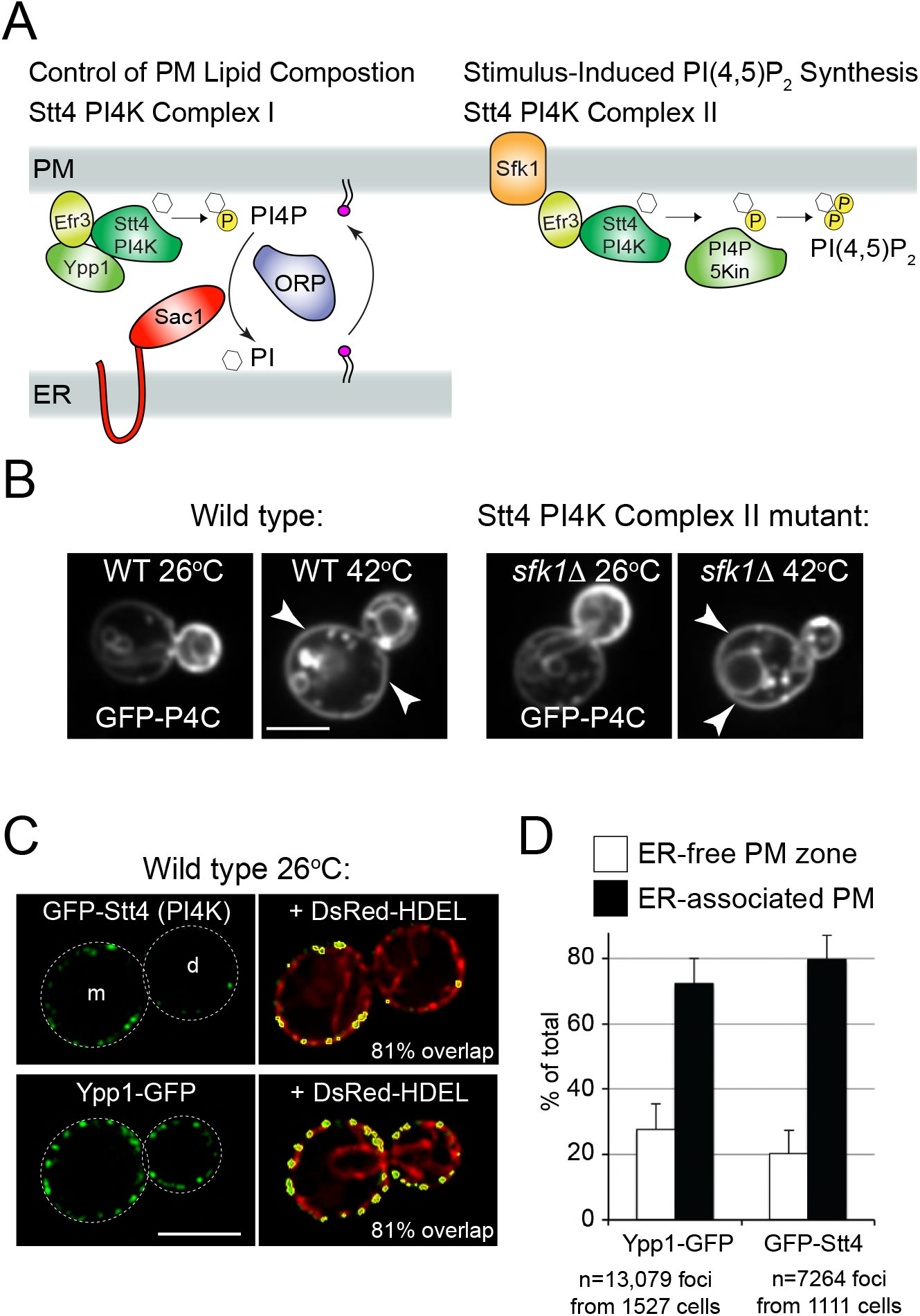
Stt4 PIK patches localize to ER-PM contact sites. **(A)** Schematic representation of two Stt4 PI 4-kinase complexes present at the pM. Stt4 complex I (Stt4/Efr3/Ypp1), also known as a PIK patch, is enriched in mother cells and is essential for ORP/Osh protein-mediated lipid transfer at ER-PM contacts. Complex II (Stt4/Efr3/Sfk1) is involved in stimulus-induced PI4P and PI(4,5)P_2_ synthesis. **(B)** Sfk1 (PI4K Complex II) is not required for heat-induced PI4P signaling in mother cells. GFP-P4C fluorescence indicating PI4P localization in wild type (left panels) and *sfk1*Δ cells (right panels) at 26°C and 42°C. Arrows point out increased PI4P levels at the PM of mother cells. Scale bar, 5 μm. **(C)** Stt4 complex I (PIK patches) are ER-associated. GFP-Stt4 or Ypp1-GFP localization in wild type cells (green) co-expressing the ER marker DsRed-HDEL (red). Mother (m) and daughter (d) cells are indicated. Stt4- and Ypp1-containing PIK patches (outlined in yellow) were automatically segmented and scored for co-localization with the ER marker. Scale bar, 4 μm. **(D)** High-content quantitative analysis of PIK patch localization with the ER. Maxima from GFP-Stt4 puncta (7264 from 1111 cells) and Ypp1-GFP puncta (13079 from 1527 cells) were identified using Fiji. Maxima positive for the ER marker DsRed-HDEL were scored as ER-associated. Results show the mean and standard deviation from three independent experiments.

### Stt4 PI4K Assemblies Localize to ER-PM Contacts

Consistent with PI4P regulation in mother cells, cortical assemblies of Stt4 PI4K complex I–also termed PIK patches–are enriched in mother cells (Audhya and Emr, 2002; Baird *et al*., 2008). It is proposed that Stt4 PIK patches may reside at junctions between the PM and endoplasmic reticulum (ER), termed ER-PM contacts, but this has not yet been demonstrated (Baird *et al*., 2008). We therefore investigated potential regulatory mechanisms for Stt4 complex I by examining PIK patch localization and interactions in more detail. We used high-content quantitative microscopy to examine PIK patch distribution in cells coexpressing a GFP-tagged PIK patch protein (Stt4 or Ypp1) and an ER marker (DsRed-HDEL). PIK patches were automatically segmented and maxima for GFP signal intensities within individual PIK patches were subsequently identified.

Next, the intensity of the DsRed-HDEL (ER) signal within each GFP maxima (PIK patch) was quantified. The dynamic range of DsRed-HDEL intensities in the ER was measured in numerous cells to define a minimal specific level for the DsRed-HDEL ER signal and maximal non-specific background DsRed-HDEL levels in ER-free regions. These measurements set a threshold value to assign individual GFP maxima as either ER-associated or ER-free.

Stt4- and Ypp1-containing PIK patches extensively coincided with the peripheral ER (Fig 2C, yellow traces show segmented GFP-Stt4 and Ypp1-GFP PIK patches; > 80% overlap with DsRed-HDEL in the examples). The DsRed-HDEL ER marker overlapped with 80% of GFP-Stt4 patches identified in high-content experiments (7264 maxima from 1111 cells in three independent experiments, Fig 2D). Likewise, 76% of Ypp1-GFP cortical patches analyzed (13,079 maxima from 1527 cells) were ER-associated (Fig 2D). Loss of the reticulon proteins (Rtn1, Rtn2, and Yop1) that shape the ER network into highly-curved tubules and membrane sheets with curved edges (West *et al*., 2011) did not alter PIK patch association with the cortical ER. GFP-Stt4 remained extensively associated with cortical ER sheets (Fig EV2B, 78% overlap), and GFP-Stt4 was notably absent from regions of the PM lacking cortical ER in the reticulon mutant cells (Fig EV2B). In contrast, localization of PIK patches with the ER depended on ER-PM contact formation. In cells lacking the ER-localized Scs2 and Scs22 proteins that form and function at ER-PM contacts (Manford *et al*., 2012), GFP-Stt4 patches were clearly observed at regions of the PM lacking cortical ER (*scs2*Δ *scs22*Δ mutant cells, Fig EV2B). Thus, Stt4 PI4K complex I assemblies (PIK patches) are associated with the cortical ER and this arrangement may impact PI4P metabolism in mother cells.

### ER-Localized Scs2 Interacts with PIK Patches at the PM and the Sac1 PI4P Phosphatase in the ER

We investigated interactions between Stt4 PI4K complex I subunits and Scs2 in further detail. Scs2, a VAP-A/B ortholog, is a tail-anchored ER membrane protein with an N-terminal cytoplasmic MSP domain. The MSP domain binds proteins with a FFAT motif (two phenylalanines in an acidic tract) (Murphy and Levine, 2016). Examination of Stt4 PI4K complex I proteins with an algorithm that scores FFAT motifs (Murphy and Levine, 2016) revealed a FFAT-like motif in the C-terminus of the Efr3 protein (…GENQN**DDFKDANED**LHSLSSRGKIFSST_782_). We then employed bimolecular fluorescence complementation (BiFC or split GFP; Fig EV3A) assays to address whether the ER-localized Scs2 protein can physically interact with the PM-localized Efr3 protein. Cortical GFP patches were observed in cells coexpressing Efr3-GFP_N_ and GFP_C_-Scs2 fusions (Fig 3A). Interestingly, the Efr3-Scs2 interaction occurred in mother cells but not daughter cells, and at cortical ER structures but not in cytoplasmic ER or nuclear ER (Fig 3A). The FFAT motif in Efr3 was necessary for efficient interaction with Scs2, as the split GFP signal intensity significantly decreased in cells expressing a truncated Efr3ΔFFAT-GFP_N_ fusion and GFP_C_-Scs2 (Figs 3A-B). In control experiments, the full-length Efr3-GFP_n_ and truncated Efr3ΔFFAT-GFP_N_ fusions interacted with Efr3-GFP_C_ equally well, indicating that the truncated form of Efr3 lacking the FFAT motif was expressed (Figs 3C-D). In addition to Scs2, Efr3 also interacted with the ER-localized PI4P phosphatase Sac1 by BiFC (Fig EV3B). Similar to Efr3, Sac1 displayed strong homotypic interactions at cortical patches (Fig EV3B), suggesting Sac1 may form oligomers in the cortical ER. In additional control experiments, Efr3 interacted with Ypp1 and Sac1 interacted with Scs2 (Fig EV3B), consistent with published data (Baird *et al*., 2008; Manford *et al*., 2012; Wu *et al*., 2014). Thus, ER-localized Scs2 interacts with the Stt4 PIK patch subunit Efr3 at the PM and the Sac1 PI4P phosphatase in the ER. This configuration may allow for the efficient control of PI4P generated by Stt4 at the PM in mother cells.

**Figure 3.**
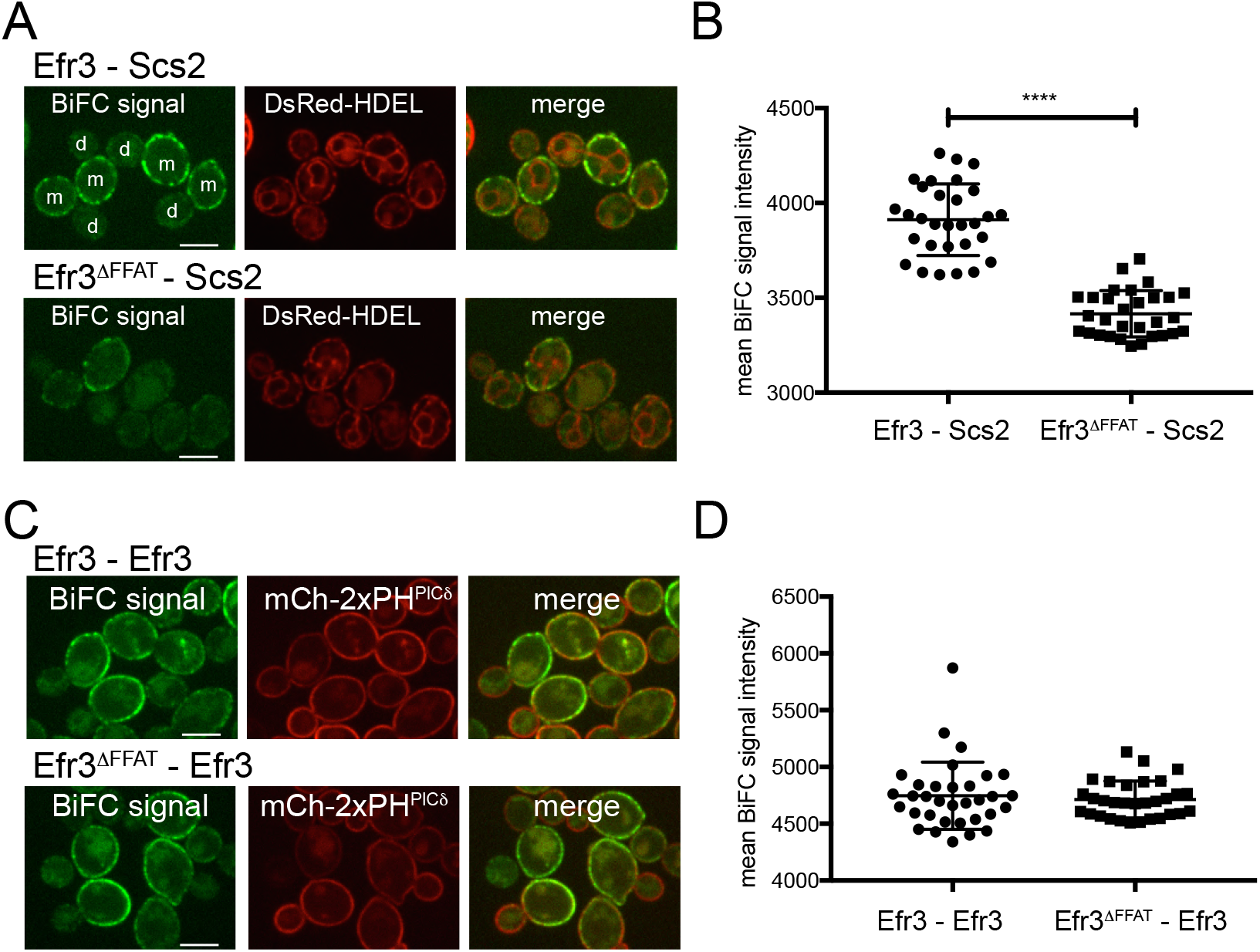
The PIK patch subunit Efr3 interacts with ER-localized Scs2. **(A)** The Efr3 FFAT motif promotes Scs2-Efr3 interactions. Interactions between Scs2 and Efr3 or Efr3ΔFFAT as scored by BiFC. Cells expressed Efr3-GFP_N_ (upper panel) or Efr3ΔFFAT -GFP_N_ (lower panel) along with GFP-Scs2 and the ER marker DsRed-HDEL. Scale bars, 5 μm. **(B)** Quantitation of specific BiFC signals generated by interaction of Efr3-GFP_N_ or Efr3ΔFFAT -GFP_N_ with GFP_C_-Scs2 at the ER using DsRed-HDEL to define regions of interest. Graph shows the mean values and standard deviations from three independent experiments. Each point represents the mean of an image frame (with >10 cells/frame); 30 frames (>300 cells) were analyzed for each. The difference between the BiFC signal in cells expressing Efr3-GFP_N_ versus Efr3ΔFFAT -GFP_N_ with GFP_C_-Scs2 was statistically significant (t test, ****p< 0.0001). **(C)** The Efr3 FFAT motif is not required for Efr3-Efr3 interactions. Cells expressed Efr3-GFP_N_ (upper panel) or Efr3ΔFFAT -GFP_N_ (lower panel) together with Efr3-GFP_C_ and the PM marker mCherry-2xPH^PLCδ^. Scale bars, 5 μm. **(D)** Quantitation of specific BiFC signals generated by interaction of Efr3-GFP_N_ or Efr3ΔCT-GFP_N_ with Efr3-GFPc using mCherry-2xPH^PLCδ^ to specify Efr3-Efr3 interactions at the PM. Graph shows the means and standard deviations from three independent experiments. Each point represents the mean value from an image frame (>10 cells/frame). Total number of frames: Efr3-Efr3 n= 33, Efr3ΔFFAT-Efr3 n= 31. The results show no difference between Efr3-GFP_N_ versus Efr3ΔFFAT -GFP_N_ with Efr3-GFPc.

### Osh3-Mediated PI4P Hydrolysis is Attenuated During Heat Shock

Our results show that Stt4 PIK patches localize to ER-PM contacts, possibly by Efr3 interactions with the ER-localized Scs2 and Sac1 proteins (Figs 2 and 3). Upon heat shock, ER-PM contacts may disassemble or Stt4 may move to ER-free PM zones, impairing Sac1-mediated PI4P hydrolysis in mother cells. However, Stt4 PIK patches remain extensively co-localized with an intact cortical ER network during heat shock (Figs EV4A-B). Thus, heat-induced PI4P signaling at the PM of mother cells is triggered by a different mechanism that we next sought to elucidate.

In addition to Efr3 and Sac1, Scs2 binds and recruits the FFAT-containing Osh3 protein (Loewen and Levine, 2005; Stefan *et al*., 2011). Osh3 is a member of a conserved family of lipid transfer proteins, the oxysterol-binding protein related proteins (ORP). The yeast Osh2 and Osh6/7 proteins transfer newly synthesized ergosterol and phosphatidylserine, respectively, from the ER in exchange for PI4P at the PM (Schulz *et al*., 2009; Moser von Filseck *et al*., 2015). As such, PI4P metabolism drives the directional transport of lipids from the ER to the PM (Fig 2A). It is currently not known what lipid(s) Osh3 transports from the ER to the PM, but Osh3 has been shown to bind PI4P and to activate the Sac1 PI4P phosphatase *in vitro* (Stefan *et al*., 2011; Tong *et al*., 2013). We reasoned that Osh proteins involved in metabolism of PM PI4P pools (by either delivering PI4P to the ER or directly to the Sac1 phosphatase) might be regulated during heat shock. Under normal growth conditions, Osh3-GFP is observed at cortical patches as well as diffusely localized in the cytoplasm (Fig 4A). Upon heat shock, Osh3-GFP PM localization (cells co-expressed the PM marker mCherry-2xPH^PLCδ^) was significantly reduced (Figs 4A-B). Instead, Osh3-GFP localized to numerous intracellular puncta following heat shock (Fig 4A).

**Figure 4.**
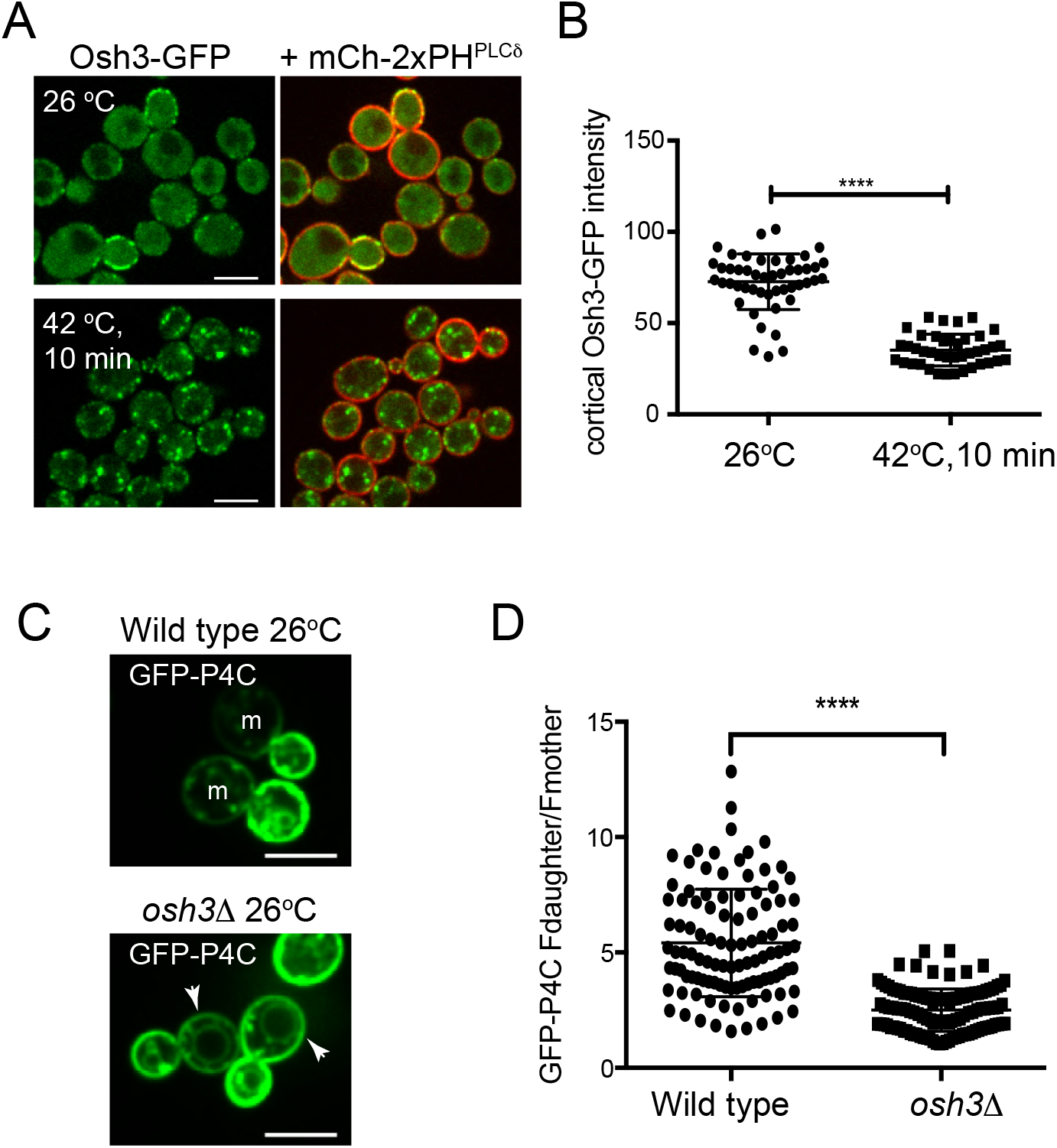
Osh3 regulates PI4P metabolism and distribution. **(A)** Localization of Osh3-GFP and the PM marker mCherry-2xPH^PLCδ^ at 26°C and after 10 min heat shock at 42°C. Upon heat shock, Osh3 switches from cortical patches to internal puncta. **(B)** Quantitation of cortical Osh3-GFP fluorescence intensity in cells at 26°C versus 10 min 42°C heat shock. The PM marker mCherry-2xPH^PLCδ^ was used to define cortical regions of interest. Graph shows the means and standard deviations from three independent experiments. Points present mean values of individual image frames (>10 cells/frame). Total number of frames analyzed: 26°C n = 48, 42°C n=41. The decrease of cortical Osh3-GFP fluorescence after heat shock was statistically significant (t test, ****p< 0.0001). **(C)** Wild type (upper panel) and *osh3*Δ□□lower panel□□cells expressing the PI4P reporter GFP-P4C grown at 26°C. Arrows point to PI4P at the PM in mother (m) cells. **(D)** Quantitation of cortical GFP-P4C fluorescence intensity in wild type and *osh3Δ*□cells grown at 26°C. See Fig 1C for details of analysis. Graph shows the Fd/Fm ratios of individual cells from three independent experiments, bars represent mean and standard deviations (t test, ****p< 0.0001). Number of cells analyzed: WT n= 105, *osh3*Δ n= 117. Scale bars, 5 μm

In control experiments, Osh2 and Osh7 remained at cortical patches upon heat shock (Fig EV4C). This suggested that mis-localization from cortical sites and thus inactivation of Osh3 may be sufficient to generate the heat-induced PI4P signal in mother cells. In support of this, the polarized distribution of PI4P was significantly reduced in *osh3*Δ cells, as GFP-P4C was increased at the PM in mother cells lacking Osh3, even under non-stress conditions (Fig 4C). Consequently, the ratio of PI4P distribution between daughter and mother cells was significantly decreased in cells lacking Osh3 (F_d_/F_m_=5.4 in wild type cells at 26°C versus F_d_/F_m_=2.5 in *osh3*Δ cells at 26°C; Fig 4D). Together, these results show that Osh3 is important for polarized PI4P distribution under non-stress conditions and is specifically attenuated in the control of heat-induced PI4P signaling.

### The Osh3 GOLD Domain Undergoes Phase Transitions

We further examined the Osh3 aggregates formed during heat shock. Remarkably, Osh3-GFP extensively localized with Hsp104 upon heat shock (Figs 5A, 5B). Hsp104 is a disaggregase that functions to refold denatured proteins upon stress conditions including heat shock (Sweeny and Shorter, 2016). Thus, Osh3 may undergo discrete phase transitions depending on environmental conditions. Under normal growth conditions Osh3 forms cortical assemblies associated with the PM, but upon heat shock Osh3 converts to intracellular granules that contain the Hsp104 chaperone (Figs 5A, 5B).

**Figure 5.**
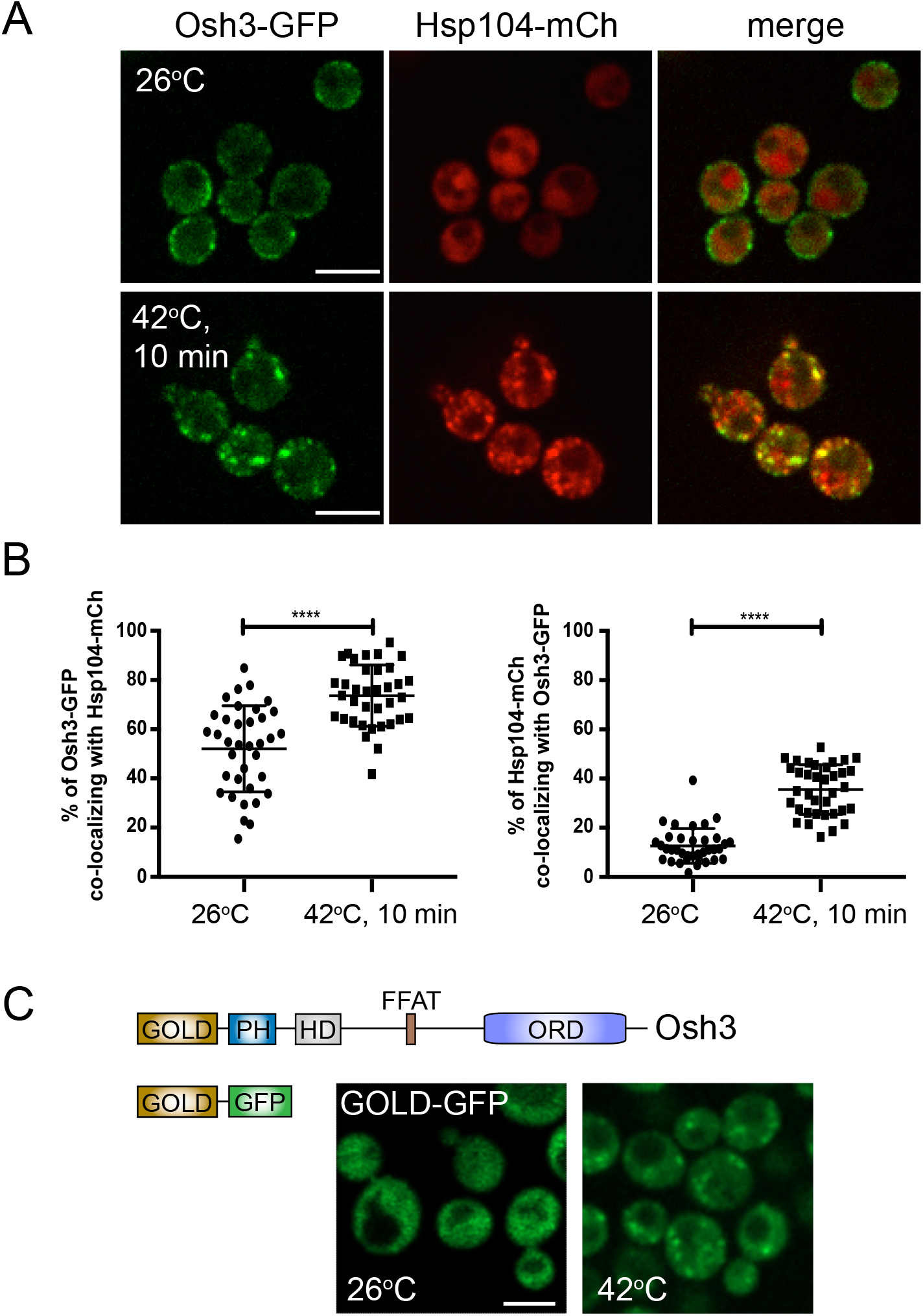
Osh3 co-localizes with Hsp104 under heat stress conditions. **(A)** Cells expressing Osh3-GFP (green) and Hsp104-mCherry (red) grown at 26°C (upper panel) and after a 10 min 42°C heat shock (lower panel). Scale bar, 5 μm. **(B)** Quantitation of Osh3-GFP and Hsp104-mCherry co-localization at 26°C and 42°C heat shock. Graphs shows the means and standard deviations from three independent experiments (t test, ****p< 0.0001). The points present values of individual image frames (>10 cells/frame). Number of frames analyzed: 26°C n = 36, 42°C n = 36. **(C)** Schematic representation of the Osh3 protein domains and the truncated GOLD^Osh3^-GFP fusion protein. Cells expressing GOLD^Osh3^-GFP at 26°C and after 10 min 42°C heat shock. Scale bar, 4 μm.

We next investigated how Osh3 stress granules form during heat shock. Surprisingly, Osh3 aggregation did not require Hsp104 or the Hsp42 proteins (Table EV1) implicated in the formation of Hsp104-containing Q-bodies (Escusa-Toret *et al*., 2013). Heat-induced Osh3 condensates also formed in the absence of Rsp5 E3 ubiquitin ligase activity and the heat-responsive protein kinases Pkc1 and Tor2 (Table EV1) (Delley and Hall, 1999; Omnus *et al*., 2016). Remarkably, Osh3 aggregation occurred independently of numerous stress-activated response factors (Table EV1). This led us to ask if some intrinsic property of the Osh3 protein itself triggers a heat-induced phase transition.

In contrast to Osh3, the Osh2 protein remains in cortical assemblies during heat shock (Fig EV4C). A distinguishing feature between these two proteins is an N-terminal GOLD domain in Osh3 that is not present in Osh2 (Fig 5C). Interestingly, the GOLD domain was sufficient for heat-induced aggregation. Under non-stress conditions, a GOLD-GFP fusion localized diffusely throughout the cytoplasm, but upon heat shock GOLD-GFP formed intracellular puncta (Fig 5C). Consistent with these *in vivo* results, the purified Osh3 N-terminal region (GST-Osh3^1-603^ including the GOLD-PH-HD-FFAT domains; Fig EV5A) displayed heat-sensitive properties *in vitro*. In sedimentation assays, GST-Osh3^1-603^ remained in the soluble supernatant fraction at 26°C, but efficiently pelleted upon incubation at high temperature (55°C, 10 min; Fig EV5B). GST-Osh3^1-603^ began to sediment upon brief physiological heat shock at 42°C (approximately 20% of the total; Fig EV5C). Heat shock is known to lower cytosolic pH (Panaretou and Piper, 1992), and so we tested whether a drop in pH (from pH 7.5 to 6.5) might contribute to Osh3 phase transitions. GST-Osh3^1-603^ sedimentation increased two-fold upon incubation at 42°C pH 6.5 (approximately 40% of the total; Fig EV5C). In control experiments, GST did not sediment at 42°C (at pH 7.5 or 6.5; Fig EV5C). Thus, the Osh3 protein displays heat and pH sensitivity *in vitro* and aggregates *in vivo* upon environmental stress conditions via its GOLD domain.

### PI4P Metabolism Controls Cell Polarity

Our results show that Osh3 controls PI4P distribution. Accordingly, the Scs2/22 proteins that recruit and regulate Osh3 at ER-PM contacts also control PI4P distribution (Manford *et al*., 2012). We therefore investigated whether defects in ER-PM contact formation result in polarity defects.

PI(4,5)P_2_ is an important spatial landmark in budding yeast. It is enriched at sites of polarized growth including incipient bud sites and mating projection tips where it recruits downstream effector proteins during polarized growth and cell signaling (He *et al*., 2007; Garrenton *et al*., 2010). As PI4P metabolism controls PI(4,5)P_2_ metabolism, alterations in ER-PM contact formation may impact polarized distribution of PI(4,5)P_2_ at the PM. To address this, we examined PI(4,5)P_2_ gradients at mating projections (‘shmoo’ tips) in wild type control cells and in *scs2 scs22 ist2* mutant cells where ER-PM contact formation is impaired >50% and PI4P levels are increased approximately two-fold (Manford *et al*., 2012). In control cells, the PI(4,5)P_2_ FLARE (GFP-2xPH^PLCδ^) was enriched at the tips of polarized mating projections formed in response to the pheromone α-factor (Fig 6A). Nearly 80% of control cells displayed enrichment of PI(4,5)P_2_ at shmoo tips (Fig 6B). In contrast, PI(4,5)P_2_ gradients were not specified to mating projections in *scs2 scs22 ist2* cells (Fig 6A). Less than 40% of the mutant cells displayed polarized enrichment of PI(4,5)P_2_ at shmoo tips (Fig 6B). Moreover, the polarized targeting of the pheromone receptor Ste2 to mating projections was impaired upon loss of ER-PM contacts (Fig EV6). Thus PI4P metabolism controls the polarized distribution of PI(4,5)P_2_ and the polarized trafficking of a cell-surface signaling receptor.

**Figure 6.**
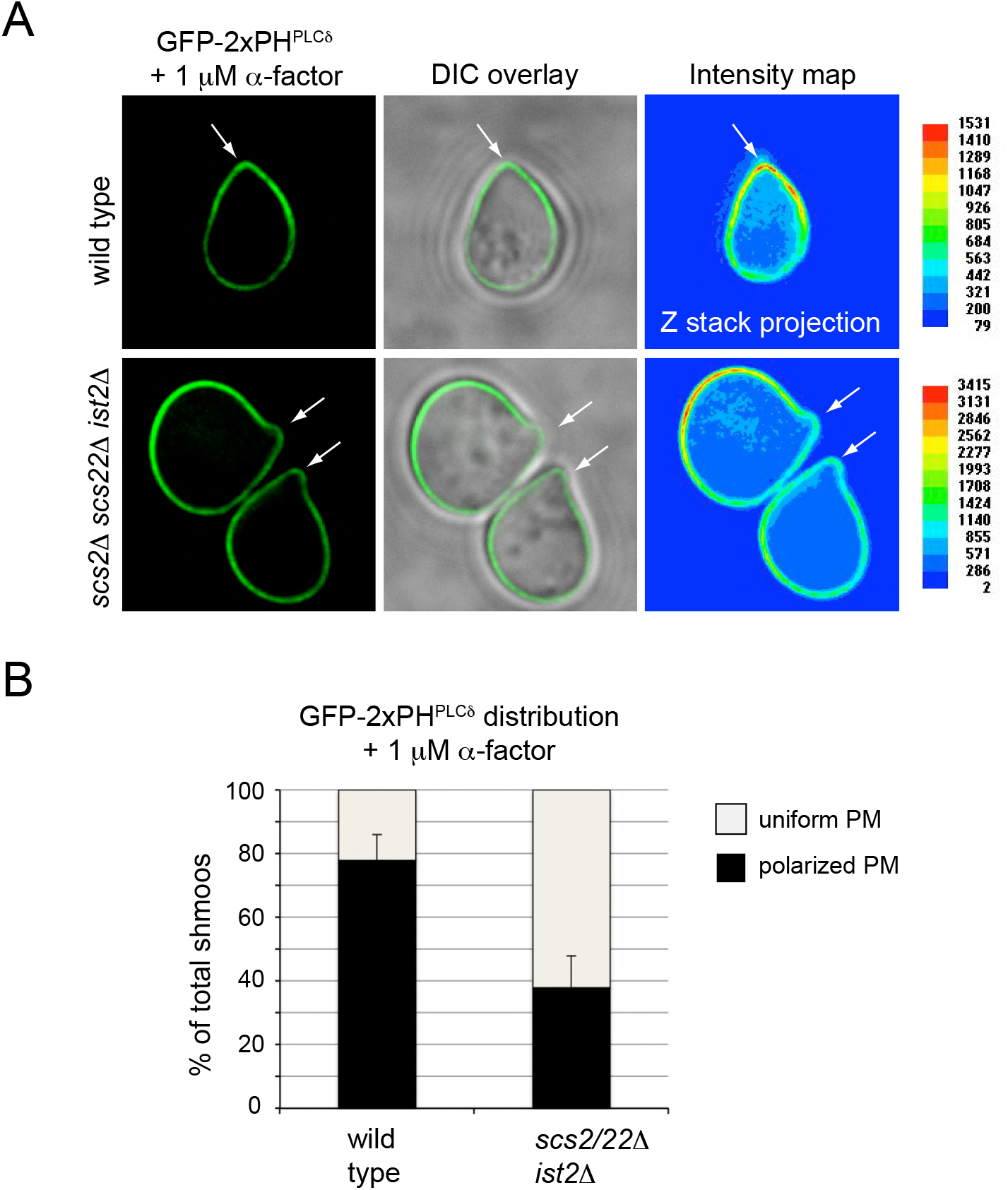
ER-PM contacts play a role in generating a plasma membrane PI(4,5)P_2_ gradient. **(A)** Exponentially growing wild type or *scs2*Δ *scs22*Δ *ist2*Δ cells expressing the PI(4,5)P_2_-binding fluorescent probe GFP-2xPH^PLCδ^ were exposed to 1 μM α-factor for 90 min before being placed on a coverslip for imaging. Representative single optical sections showing GFP-2xPH^PLCδ^ localization in wild type and mutant cells are provided. The false color image represents the indicated scale of measured pixel intensity in a Z-stack projection. Arrows point to tips of mating projections. **(B)** Quantitation of PI(4,5)P_2_ gradient formation in wild type (n=55) and *scs2*Δ *scs22*Δ *ist2*Δ (n=51) cells. Graph shows the means and standard deviations from three independent experiments.

### Stimulus-Induced Regulation of ORP5 Localization

We wondered if other ORP family members undergo dynamic changes in response to external stimuli, similar to the Osh3 protein. The ORP5 protein localizes to ER-PM contacts where it carries out phosphatidylserine-PI4P exchange reactions (Chung *et al*., 2015b; Sohn *et al*., 2016; Sohn *et al*., 2018). ORP5 cortical localization is mediated by a PH domain that binds PI(4,5)P_2_ in the PM (Ghai *et al*., 2017; Sohn *et al*., 2018). Thus, ORP5 localization may be regulated in response to stimuli that trigger phospholipase C (PLC) activation and PI(4,5)P_2_ hydrolysis. We used the calcium (Ca^2+^) ionophore ionomycin known to result in intense PLC activity and significant reductions in PI(4,5)P_2_ levels at the PM (Varnai and Balla, 1998; Idevall-Hagren *et al*., 2015). As monitored by total internal reflection fluorescence (TIRF) microscopy in HeLa cells, addition of 10 μM ionomycin (that lowers PI(4,5)P_2_ levels) resulted in the rapid loss of GFP-ORP5 cortical localization (Fig EV7A). This ORP5 regulatory mechanism may simply work through dynamic fluctuations in PH domain-PI(4,5)P_2_ interactions (Ghai *et al*., 2017; Sohn *et al*., 2018), as ORP5 does not have a GOLD domain. But importantly, loss of ORP5/8 is known to result in increased PI4P and PI(4,5)P_2_ levels at the PM (Chung *et al*., 2015b; Sohn *et al*., 2016; Ghai *et al*., 2017; Sohn *et al*., 2018). Thus, attenuation of ORP-mediated phosphoinositide hydrolysis may be a conserved mechanism for the rapid generation of PI4P and PI(4,5)P_2_ at the PM in response to physiological stimuli.

## Discussion

Our results suggest PI4P metabolism is a key determinant in the control of polarized growth. Importantly, PI4P distribution can be rapidly modulated at the plasma membrane in response extracellular signals to direct polarity cues as needed.

### Phosphoinositide Metabolism Controls Polarized Cell Growth

PI4P is enriched in the plasma membrane of rapidly growing small-budded yeast cells (during late G1 and S) (Fig 1). Likewise, PI(4,5)P_2_ is also enriched at sites of polarized growth in yeast (incipient bud sites, ‘shmoo’ tips, and cleavage furrows) (Garrenton *et al*., 2010). How PI4P and PI(4,5)P_2_ gradients are established is not well understood. Surprisingly, the Stt4 PI4K that synthesize PI4P at the PM is found primarily in mother cells and is not readily apparent in small-budded daughter cells where PI4P is enriched (Audhya and Emr, 2002; Baird *et al*., 2008) (Fig 2). This paradoxical difference in the distribution of the Stt4 PI4K and its product PI4P may be explained if PI4P pools generated in mother cells are rapidly turned over. Consistent with this notion, Stt4 PIK patches localize extensively to regions of the plasma membrane (PM) associated with the cortical endoplasmic reticulum (ER) containing the PI4P phosphatase Sac1 (Figs 2, EV2, 3, EV3).

Our data suggest that Stt4 localization is mediated through interactions between the PIK patch component Efr3 and the ER-localized VAP orthologs Scs2/22. Loss of the Scs2/22 proteins does not disrupt Stt4 PIK patch formation at the PM, but does impair PIK patch ER-association (Fig EV2). Accordingly, deletion of the FFAT motif in Efr3 reduced interaction with Scs2 (Fig 3). To our knowledge, our study provides the first evidence that a PI4K, the yeast PI4KII Iα ortholog Stt4, resides at ER-PM contacts. It is unclear if mammalian PI4KIIIα localizes to ER-PM contacts, but phosphatidylinositol transfer proteins that regulate PI4KIIIα activity function at ER-PM contacts (Chang *et al*., 2013; Kim *et al*., 2015). PI4P turnover also depends upon the ER-localized Scs2/22 proteins and the Sac1 phosphatase (Foti *et al*., 2001; Stefan *et al*., 2011; Manford *et al*., 2012). Thus, both PI4P synthesis and turnover occurs at ER-PM contacts.

Placing PI4KIIIα at ER-PM junctions does not necessarily mean that PI4P synthesis and degradation occur in a futile cycle at ER-PM contacts. Rather, localization of the Stt4 PI4K at ER-PM contacts provides insight into the regulation and function of PI4P at these cellular structures (Fig 7A). In addition to Efr3, the Scs2/22 VAP proteins bind and recruit the Osh2 and Osh3 proteins to ER-PM contacts (Loewen and Levine, 2005; Stefan *et al*., 2011). Osh2 and Osh3 are members of the ORP lipid transfer protein family that deliver newly synthesized lipids from the ER to the PM in exchange for PI4P (Fig 7A). Osh2 is proposed to transfer sterol lipids from the ER to the PM (Schulz *et al*., 2009). It is not yet known what lipid species are transferred by the Osh3 protein, but Osh3 can stimulate PI4P hydrolysis by the Sac1 phosphatase *in vitro* (Stefan *et al*., 2011). Interestingly, the Scs2 VAP ortholog also interacts with the Sac1 phosphatase in the ER (Fig EV3) (Manford *et al*., 2012). Thus by positioning both PI 4-kinase and PI4P phosphatase activities at ER-PM contacts, the Osh2 and Osh3 proteins may be able to execute multiple rounds of non-vesicular lipid exchange reactions at a single site.

**Figure 7.**
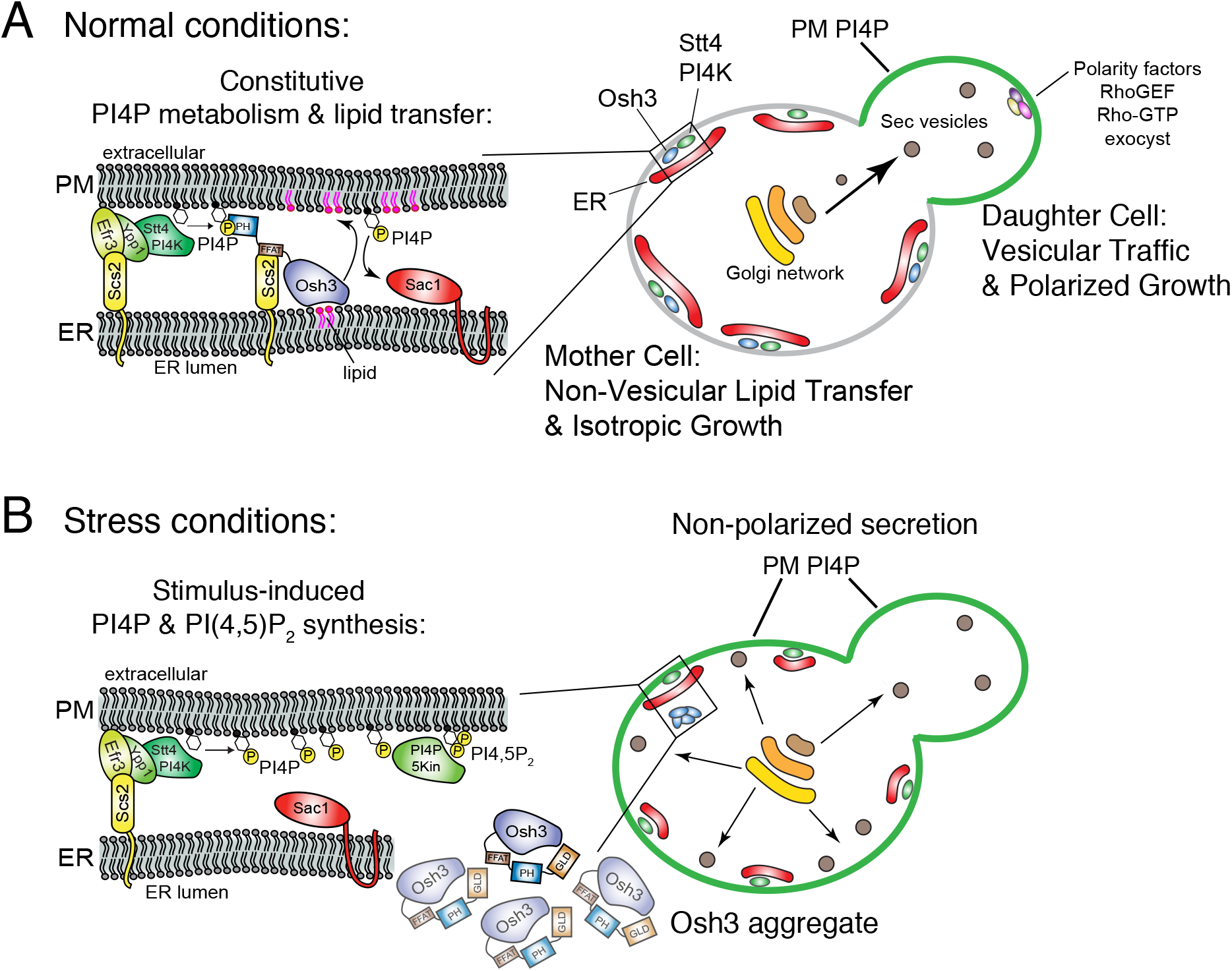
Model for PI4P regulation during polarized growth and stress conditions. **(A)** Under normal growth conditions, PI4P levels are regulated at ER-PM contact sites through its production by Stt4 PIK patches and turnover by Osh3 and Sac1. Osh3-mediated PI4P metabolism is coupled to exchange for an as yet unknown lipid at the PM. PI4P levels at the PM are higher in the growing daughter cell (bud) than in the mother cell, likely due to the lack of an established cortical ER network in the newly formed bud. As a consequence, PI4P- and PI(4,5)P_2_-regulated vesicle trafficking is directed to the growing daughter cell. **(B)** Under heat shock conditions, Osh3 localization and function at ER-PM contacts is lost as Osh3 forms cytoplasmic stress granules. Diminished Osh3-mediated PI4P turnover contributes to increases in PI4P and possibly PI(4,5)P_2_ levels at the PM. Loss of PI4P and PI(4,5)P_2_ gradients at the PM triggers isotropic secretion and cell signaling cascades necessary to maintain PM integrity.

Importantly, PI4P serves at least two vital roles in membrane lipid dynamics: non-vesicular lipid transport and vesicular trafficking. In mother cells with an extensive cortical ER network (West *et al*., 2011), PI4P is utilized as a currency for ORP-mediated lipid exchange. Non-vesicular exchange of PI4P for another lipid does not result in a net lipid gain at the PM and may not drive rapid membrane expansion. However, ORP-mediated exchange reactions may be critical for proper PM lipid composition and organization, thus ensuring PM integrity in mother cells while polarized vesicular trafficking is directed to the growing bud (during late G1/S) (McCusker and Kellogg, 2012). ORP-mediated exchange reactions would also ensure that PI4P levels are kept low in mother cells, resulting in the polarized distribution of PI4P between mother and small-budded daughter cells. It is estimated that nearly 90% of Stt4-generated PI4P is rapidly consumed by the Sac1 phosphatase (Foti *et al*., 2001); the bulk of this PI4P turnover may take place in mother cells via ORP-mediated lipid exchange.

In contrast, PI4P is readily available at the PM of rapidly growing small-budded daughter cells. PI4P may serve as a spatial landmark along with PI(4,5)P_2_ and other anionic lipids for the targeting of polarity factors including Rho-family small GTPases, their associated guanine nucleotide exchange factors, and exocyst subunits that determine sites for the polarized targeting of secretory vesicles (Audhya and Emr, 2002; He *et al*., 2007; Fairn *et al*., 2011).

Consistent with this idea, both PI4P and PI(4,5)P_2_ suffice for targeting polybasic proteins to the PM in mammalian cells (Hammond *et al*., 2012). While the ER is inherited in newly formed daughter cells, polarized trafficking of secretory vesicles continues and drives PM expansion in the growing bud until late G2/M, at which point the daughter cell establishes an extensive cortical ER network and switches from polarized to isotropic growth (West *et al*., 2011; McCusker and Kellogg, 2012). The lack of an extensive cortical ER network (and the Sac1 phosphatase) may explain how PI4P accumulates in small-budded daughter cells that seem to be devoid of Stt4 PIK patches and even contain cortical Osh3 assemblies.

In simple terms, the PM may be considered as ER-associated and ER-free. Non-vesicular membrane lipid transfer events take place at ER-associated PM zones (predominately in mother cells), while vesicular membrane trafficking events take place in ER-free PM zones (*i.e*. polarized secretory trafficking to the bud) (Fig 7). ER-PM contacts are proposed to act as a physical barrier preventing exocytosis (Ng *et al*., 2018). This is reasonable, as ER-PM contacts are closely apposed (10-30 nm apart) and may not accommodate secretory vesicles (50-100 nm in diameter). However, less than 50% of the PM is ER-associated in budding yeast and the cortical ER network is constantly reorganized (Manford *et al*., 2012). In some mammalian tissue culture cells, only 2% of the PM is ER-associated (Giordano *et al*., 2013) and it is unclear how the ER could serve as an effective fence to preclude vesicle docking and fusion at the PM.

We propose that control of PI4P metabolism and distribution by ORP-mediated lipid exchange determines where sites of exocytosis occur. Upon heat shock, budding yeast cells rapidly halt polarized secretion to daughter cells and switch to isotropic trafficking between mother and daughter cells (Delley and Hall, 1999). This coincides with a rapid increase in PI4P availability in mother cells (Figs 1, EV1), possibly for heat-induced PI(4,5)P_2_ synthesis (Audhya and Emr, 2002). The Stt4 PI4K remains ER-associated upon heat shock (Fig EV4).

Instead, Osh3 cortical localization decreases (Figs 4 and 5), suggesting that attenuation of Osh3 contributes to generation of the PI4P signal in mother cells.

In support of this, loss of Osh3 increases PI4P availability in mother cells (Fig 4). A similar regulatory mechanism takes place in mammalian cells. Depletion of ORP5 and ORP8 results in increased PI4P and PI(4,5)P_2_ levels (Chung *et al*., 2015b; Sohn *et al*., 2016; Ghai *et al*., 2017; Sohn *et al*., 2018). Our observations and another recent study (Sohn *et al*., 2018) suggest that ORP5/8 activity is tuned according to changes in PI(4,5)P_2_ levels. In response to stimuli that trigger PI(4,5)P_2_ consumption, for example Ca^2+^-regulated phospholipase C activity (Varnai and Balla, 1998), ORP5 cortical localization is lost (Fig EV7A) likely due to reduced interactions between the ORP5 PH domain and PI(4,5)P_2_ at the PM (Sohn *et al*., 2018). Transient loss of ORP5/8 activity would result in a rapid increase in PI4P for PI(4,5)P_2_ re-synthesis (Fig E7B). As PI(4,5)P_2_ is regenerated, ORP5/8-mediated PI4P consumption would resume and this may explain why cells have two PI4KIIIα complexes (I and II). While complex I (PI4K-Efr3-Ypp1/TTC7) may provide an initial PI4P pool, activation of complex II (PI4K-Efr3-Sfk1/TMEM150) may ensure rapid stimulus-induced PI(4,5)P_2_ synthesis (Fig EV7B) (Audhya and Emr, 2002; Chung *et al*., 2015a).

The ability of ORP5/8 to serve as a PI(4,5)P_2_ sensor and modulator was recently described as a ‘rheostat’ mechanism for PI(4,5)P_2_ homeostasis (Sohn *et al*., 2018). This ‘rheostat’ follows Le Chatelier’s Principle that states when an external stress alters a reaction in equilibrium, the reaction will shift to restore equilibrium. Thus if PI(4,5)P_2_ levels drop, PI(4,5)P_2_ synthesis increases to maintain homeostasis. Our findings indicate that cells do not merely respond to changes in product and reactant concentrations, here PI(4,5)P_2_ and PI4P.

Rather, Le Chatelier’s Principle can be applied in response to an external stress, heat-induced in our study, to establish a new equilibrium. Heat shock attenuates Osh3 function effectively increasing PI4P. As the balance between PI and PI4P cannot be restored, a new equilibrium between PI4P and PI(4,5)P_2_ is established with levels of both increased. Upon return to non-stress conditions, restoration of Osh3 function would return the system to the resting state and re-establish the polarized distribution of PI4P. Thus the same governing principle not only allows cells to maintain PI4P and PI(4,5)P_2_ homeostasis, but also to respond to external signals by effectively adjusting PI4P and PI(4,5)P_2_ levels as needed.

### The Osh3 Protein Undergoes Phase Transitions

Control of PI4P metabolism ensures cellular homeostasis and modulates responses to extracellular stimuli, such as a change in environmental conditions.

We have found that Osh3 undergoes a phase transition in response to heat shock providing a rapid mechanism for the generation of a PI4P signal at the PM under these conditions. Upon heat shock, Osh3 shifts from cortical assemblies to intracellular granules containing the Hsp104 disaggregase (Fig 5A). However, not all Hsp104 puncta contained Osh3, suggesting Osh3 may form a distinct subset of heat-sensitive aggregates (Fig 5B). A growing number of proteins are known to undergo phase separations (*e.g*. liquid-liquid, liquid-gel, and aggregation) in response to various stress conditions including heat stress and nutrient starvation. In budding yeast, these include the Whi3 protein upon prolonged pheromone exposure, the AMPK/Snf1 regulator Std1 in response to carbon source, and the Sup35 protein upon glucose starvation (Schlissel *et al*., 2017; Simpson-Lavy *et al*., 2017; Franzmann *et al*., 2018). Thus, it is increasingly clear that protein phase separations and protein aggregation have important physiological roles during cellular stress responses.

The Osh3 protein does not appear to contain hallmarks such as a glutamine-rich region known to influence protein aggregation. However it does possess differentially charged domains that may render it sensitive to changes in pH (Franzmann *et al*., 2018). The N-terminal GOLD and C-terminal ORD regions are quite basic (pI=9.5 and 9.1, respectively) while the region spanning the FFAT motif is negatively charged (pI=3.6). The PH and helical domains have pI values near physiological cytosolic pH under non-stress conditions (pI= 7.1 and 7.2, respectively). Importantly, cytosolic pH decreases upon heat shock as the PM H+-ATPase Pma1 is a heat-sensitive protein and the Pma1 inhibitor, Hsp30, is induced by heat (Panaretou and Piper, 1992; Zhao *et al*., 2013). A reduction in cytosolic pH upon heat shock may induce Osh3 aggregation through inter-molecular electrostatic interactions between the negatively charged FFAT region and positively charged regions (the PH and HD regions would become positively charged as cytoplasmic pH drops). Consistent with this notion, the N-terminal half of the Osh3 containing the GOLD-PH-HD-FFAT domains sediments *in vitro* upon incubation at 42°C at pH=6.5 (Fig EV5).

The GOLD domain of Osh3 was sufficient for stress granule formation *in vivo* (Fig 5). Structures of the GOLD domain from other proteins including p24 family members have revealed a conserved β-sandwich fold (Nagae *et al*., 2016; McPhail *et al*., 2017). Perhaps, heat and pH fluctuations could induce rearrangements of the β-strands altering GOLD domain oligomerization, akin to β-strand re-arrangements that occur in prion-like proteins. The GOLD p24 family proteins form homo- and hetero-oligomeric complexes for the transport of cargo proteins from the ER. As pH in the ER lumen is affected by changes in cytosolic pH (Wolf *et al*., 2014), p24 subunit assembly and folding may be regulated by stress-induced pH changes. Previously, we reported that heat-induced ceramide synthesis in the ER is regulated by PI4P and PI(4,5)P_2_ metabolism (Omnus *et al*., 2016). Inactivation of Osh3 in the cytoplasm during heat shock may trigger ceramide- and p24-mediated protein quality control in the ER. In this manner, stress-sensitive GOLD proteins may simultaneously coordinate cytoplasmic and secretory stress responses. Intriguingly, mammalian cells possess two cytoplasmic GOLD-containing proteins, ACBD3 and SEC14L1 (Anantharaman and Aravind, 2002). ACBD3 is an integral component of a PI4K complex that synthesizes PI4P at late Golgi compartments (Klima *et al*., 2016; McPhail *et al*., 2017). It will be important to investigate whether GOLD-mediated phase transitions modulate PI4P metabolism and trafficking at Golgi compartments in response to changes in physiological and environmental conditions.

## Materials and Methods

Additional details are provided in the in the Expanded View.

### Strains and Plasmids

Yeast strains and plasmids used in this study are listed in the Expanded View. Standard techniques were used for yeast and bacterial growth. HeLa cells were cultured in Dulbecco’s modified essential Eagle medium (Life Technologies) supplemented with fetal bovine serum (Life Technologies) at 37°C and 5% CO_2_. Transfection of plasmids was carried out with Lipofectamine 3000 (Life Technologies).

### Fluorescence Microscopy

Images were obtained with a DeltaVision RT microscopy system (Applied Precision), an Ultraview Vox spinning disk confocal microscopy system (Perkin Elmer), or a Ti-E inverted microscope equipped with a TIRF illuminator (Nikon). See the Expanded View Materials and Methods for additional microscopy details.

### Quantitative Image Analysis

Quantitative image analysis was conducted using Fiji. To measure PI4P intensities at the PM of mother and daughter cells, cells expressing GFP-P4C and the PM marker mCherry-2xPH^PLCδ^ were analyzed at 26°C and after a 10 min 42°C heat shock. GFP-P4C intensity peaks co-localized with mCherry-2xPH^PLC δ□^ in mother and daughter cells were measured using line scans and used to calculate Fd/Fm ratios.

To determine PIK patch-ER association by high-content imaging, the Find Maxima tool in Fiji was used to identify GFP maxima in PIK patches and the intensity of the DsRed-HDEL ER signal for each GFP maxima was measured to assign individual GFP maxima as either ER-associated or lacking ER.

Osh3-GFP cortical intensity was measured by co-expressing mCherry-2xPH^δ^ as a PM marker. For co-localization of Osh3-GFP and Hsp104-mCherry, the percent of pixel area with GFP fluorescence that overlapped with mCherry signal and vice versa was determined.

### Recombinant Protein Purification and Sedimentation Assays

GST or GST-Osh3^1-603^ fusion proteins were purified from C41(DE3) cells with glutathione sepharose 4B (GE Healthcare). Purified GST or GST-Osh3^1-603^ were incubated at the indicated temperature and pH for 10 minutes, and centrifuged at 50,000 rpm for 20 minutes at 4°C. The resulting supernatant and pellet fractions were prepared for SDS-PAGE analysis and Coomassie-stained to detect proteins.

## Acknowledgements

We thank Scott Emr, Yuxin Mao, and Mizushima Noboru for strains and plasmids. We also thank Helen Yuan, Dan Baird and Taki Nishimura for assistance with reagents and experiments. We are grateful to Janos Kriston-Vizi, Ricardo Henriques, Tim Levine, and members of the Stefan lab for helpful discussions. The Stefan lab is supported by MRC funding to the MRC LMCB University Unit at UCL, award code MC_UU_12018/6. DJO was supported by a Wenner-Gren Foundations fellowship.

## Author Contributions

DJO, AC, GHC, JMB, and CJS designed and performed experiments and analyses. CJS directed the research and wrote the manuscript.

